# Identification of stable reference genes for quantitative PCR in koalas

**DOI:** 10.1101/215681

**Authors:** N. Sarker, J. Fabijan, R.D. Emes, F. Hemmatzadeh, J. Meers, J. Moreton, H. Owen, J. M. Seddon, G. Simmons, N. Speight, D. Trott, L. Woolford, R.E Tarlinton

## Abstract

To better understand host and immune response to diseases, gene expression studies require identification of reference genes with stable expression for accurate normalisation. This study describes the selection and testing of reference genes with stable expression profiles in koala lymph node tissues across two genetically distinct koala populations. From the 25 most stable genes identified in transcriptome analysis, 11 genes were selected for verification using reverse transcription quantitative PCR, in addition to the commonly used *ACTB* and *GAPDH* genes. The expression data were analysed using stable genes statistical software - geNorm, BestKeeper, NormFinder, the comparative ΔCt method and RefFinder. All 13 genes showed relative stability in expression in koala lymph node tissues, however *Tmem97* and *Hmg20a* were identified as the most stable genes across the two koala populations.

## INTRODUCTION

The ability to measure variation in gene expression due to host and pathogen differences, between and within individuals, and over time, is crucial for identifying genes involved in disease progression. The fluorescence based reverse transcription quantitative polymerase chain reaction (RT-qPCR) is the gold standard procedure for quantifying gene expression. In recent years, it has emerged as a vigorous and widely used technique in quantitative data analysis due to its high sensitivity, specificity, reliability, reproducibility, swiftness, ease of process and high data throughput in comparison to other quantification procedures such as microarray, northern blotting or ribonuclease protection analysis^1-4^. There is no disagreement that RT-qPCR is a robust technique for quantifying gene expression, but the reliability of RT-qPCR results depends on multiple factors such as RNA integrity and quantity, accurate reverse transcription, primer efficiency and most importantly, suitable stable internal gene selection for normalization^5^. To minimize inaccuracies in quantification of expression of genes of interest, a stable reference (housekeeping) gene is required as an endogenous control to normalize the technical variation within the experimental conditions^6^. A reference gene with unstable expression may generate misleading results and inaccurate conclusions.

Historically, many RT-qPCR studies in humans and various animal species have been normalised using the reference genes Glyceraldehyde-3-phosphate dehydrogenase (*GAPDH*) and Beta-actin (*ACTB*), however these genes can have variable expression stabilities across tissue types and experimental conditions^7,8^. Ideally, the expression level of a reference gene should remain stable across various development stages^9-11^, types of tissues^12-14^, with cancer progression^15^, under the influence of physiological hormones^16^ and under different environmental and health conditions^17,18^. However, there is no single reference gene available that remains stable in all conditions. Therefore, experimental validation of reference genes should be carried out for each type of tissue, disease state and for other relevant variables. Recently, guidelines for the minimum information required for the publication of quantitative real-time PCR experiments (MIQE) have been developed to evaluate the acceptability of RT-qPCR data, including the requirements for standardizing experiments^19^.

This study was part of a larger-scale project investigating the pathogenesis of koala retrovirus (KoRV). The koala (*Phascolarctos cinereus*) is an arboreal herbivorous marsupial species and a popular icon of Australia. The population is under threat from multiple factors to the extent that koalas are now nationally listed as vulnerable to extinction^20^. Habitat loss, dog attacks and disease are key drivers of koala population decline with particular threats being koala retrovirus (KoRV) and Chlamydiosis^21^. Lymphoid neoplasia has long been recognised as a common malignancy of koalas, and because retroviruses are known to cause neoplasia in other vertebrate species^22^, it has been hypothesised that KoRV plays a role in this disease in koalas^23-25^. Moreover, since retroviruses are also associated with immunodeficiency in their hosts, it is speculated that KoRV may be involved in the susceptibility of koalas to opportunistic infections such as chlamydiosis^26^. Across their distribution, koala populations differ in their KoRV proviral and viral RNA loads and health status, with the incidence of chlamydiosis and malignancy being higher in northern populations of koalas compared to the southern populations^21,27-29^.

A greater understanding of the association between KoRV gene expression and disease is needed. One of the barriers to measuring KoRV gene expression has been the lack of suitable reference genes for normalisation of expression values in RT-qPCR experiments. To the best of our knowledge, there are no published data available on stable expression analyses of reference genes in koala tissues. The objective of this study was to identify stable reference genes in lymph node tissue in koalas from a northern and a southern koala population.

## RESULTS

### Assessment of RNA quality

Total RNA was extracted from lymph node tissues of 19 koalas. The A260/280 ratio of all purified RNA samples was between 2.0 and 2.2 and A260/230 ratio values of all samples were greater than 2.0 (Table S1).

### Verification of primer specificity and PCR efficiency for qPCR

The primer pair specificity of candidate reference genes was verified through melt curve analysis and 2 % agarose gel electrophoresis for the respective genes’ qPCR assays. Based on qPCR efficiencies (E) value, 13 reference genes (*Bet1, Pdap1, Sec22b, Hmg20a, Ndufaf3, Smap2, Chchd2, Grk2, Nckap1l, Stx12, Tmem97, ACTB* and *GAPDH*) were selected for further analysis. Information for the primers of candidate reference genes is listed in Table 1. Gel electrophoresis confirmed that all selected gene-specific primers produced a single band at the expected size with no primer dimer or non-specific bands (Figure S1). Similarly, a single peak was observed in the melt curve analysis for each genes’ qPCR assay (Figure S2). Sequence of amplicons from 8 of the newly designed primer pairs confirmed specific amplification of the targeted genes; no usable sequence was obtained from three amplicons (*Ndufaf3, Bet1, Grk2*).

**Table 1:**
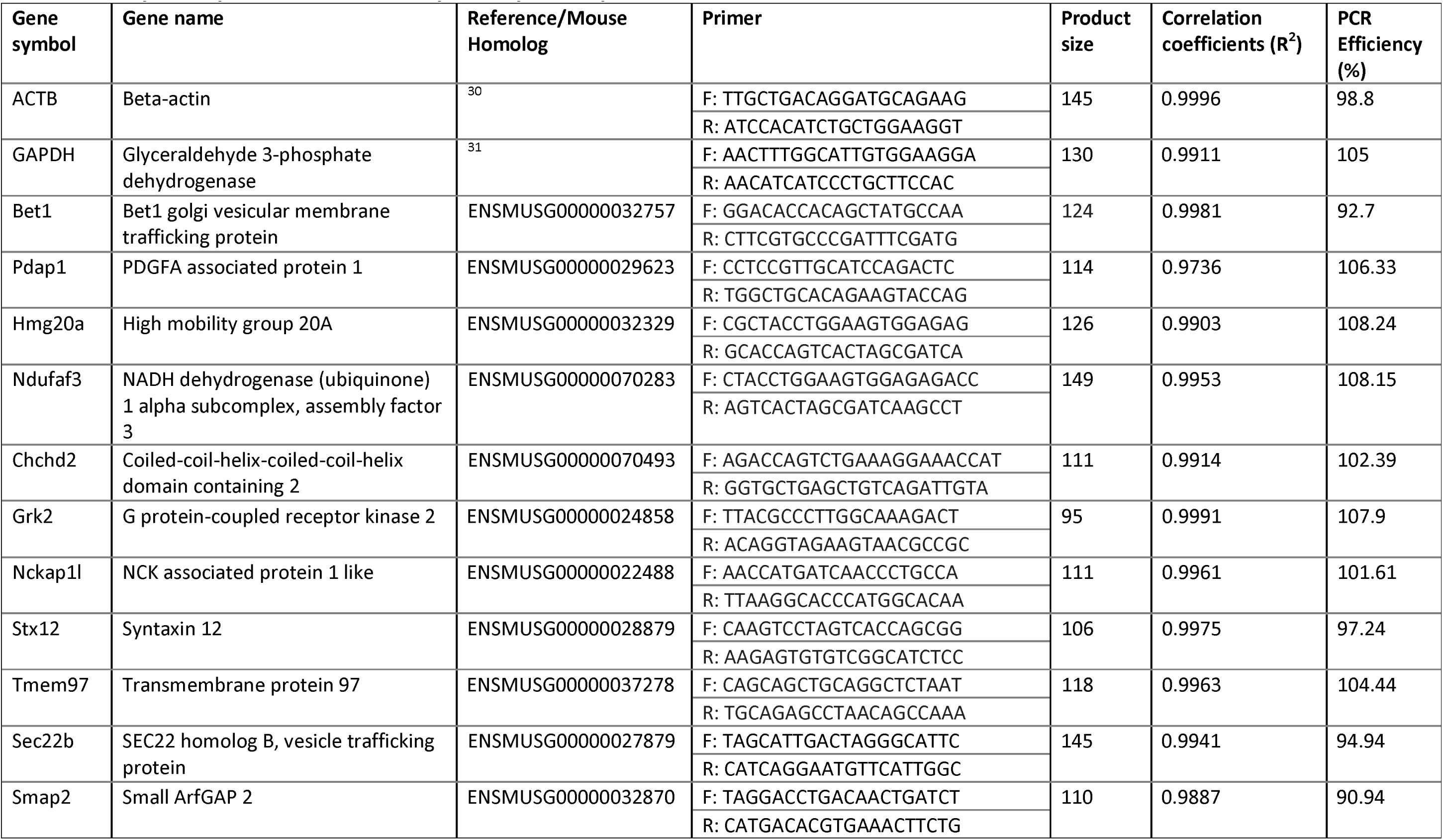
Description of primers used in this study for RT-qPCR analysis.

### Expression profile of the candidate reference genes

The steadiness of mRNA expression for each of the 13 candidate reference genes in koala lymph node tissues was analysed through Ct values. In regards to expression level, only *ACTB* was highly abundant with its average Ct values ranging from 13.17 to 15.85 (Fig. 1). Ten other candidate reference genes expressed at a medium level, with average Ct values ranging from 20.98 to 25.41, whilst *Smap* and *Sec22b* showed very low levels of expression (mean Ct= 30.27 and 29.25). The descriptive statistics of the Ct values across all the samples of each gene are available in supplementary information (Table S2).

**Figure 1:**
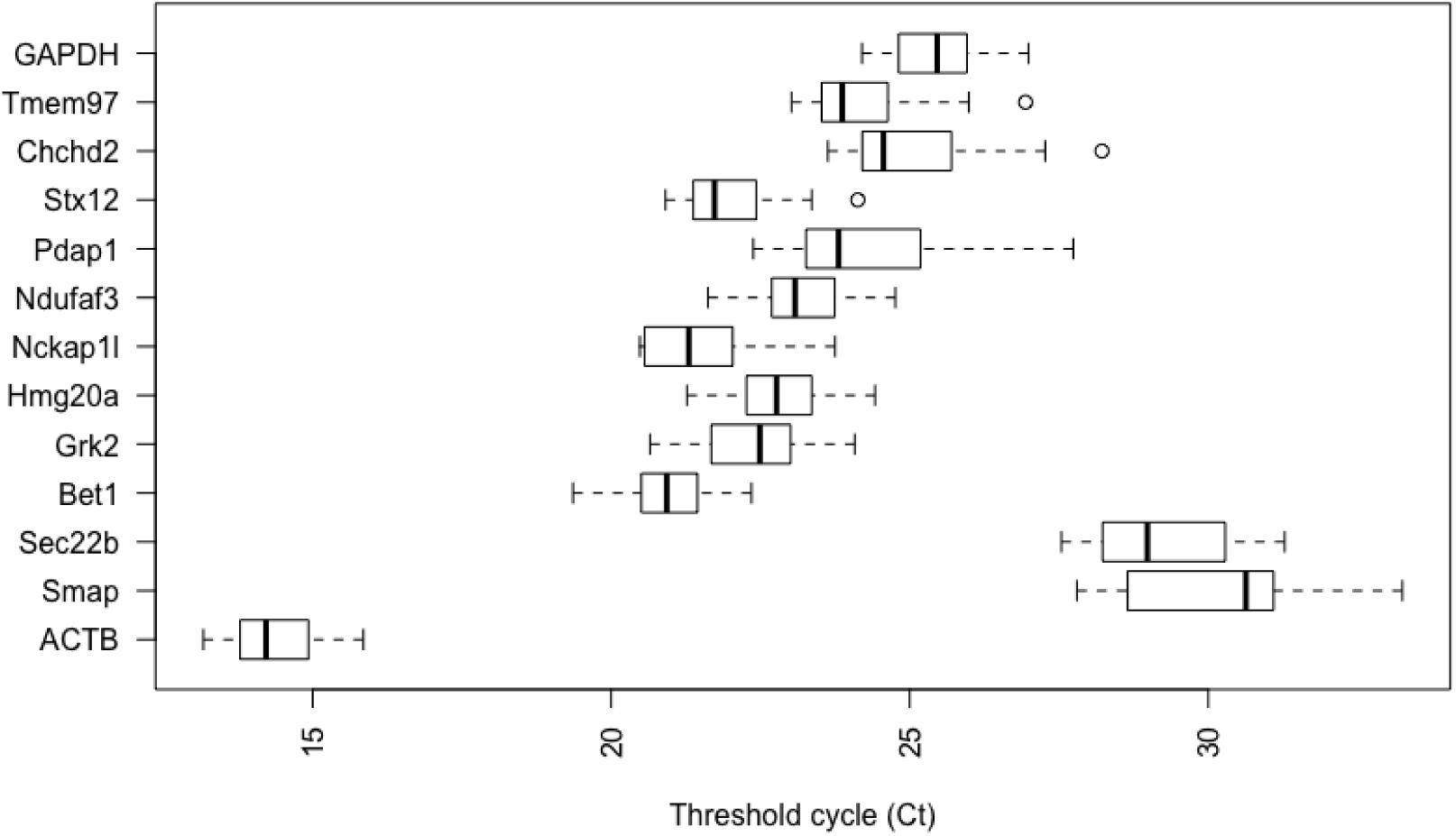
Threshold cycle (Ct) values for 13 analyzed genes obtained using qPCR in lymph node tissues of QLD and SA koalas. Box shows the 25/75 percentile; A line across the box indicates the median; Whiskers extend 1.5 times the interquartile range from the 25/75 percentiles; outliers are represented by dots; n = 19 sample points.

### Expression Stability of Candidate Reference Genes

The expression stability of 13 candidate reference genes was evaluated through geNorm, BestKeeper, NormFinder, comparative ΔCt and finally, overall stabilities were ranked using comprehensive RefFinder tool across all the QLD, SA and combined koala tissues.

#### geNorm analysis

The geNorm algorithm evaluated the stability of reference genes based on expression stability value (M), as shown in Fig. 2 A-C. All evaluated housekeeping genes had an M value below 1.5 which is the recommended geNorm cut off value for stable gene selection through RT-qPCR analysis^8^. This result confirms that the candidate reference genes were stable across lymph node tissues from different koalas. With the lowest M values, *Grk2* and *Hmg20a* were the most stably expressed genes among SA koalas whereas *Hmg20a* and *Ndufaf3* were the most stable genes in QLD koalas and also when the groups were combined. Overall, *Hmg20a* was found to be the most stable gene and *Pdap1* was the least stable gene with the highest M value in separate QLD and SA population evaluations and also in combination group analysis.

**Figure 2:**
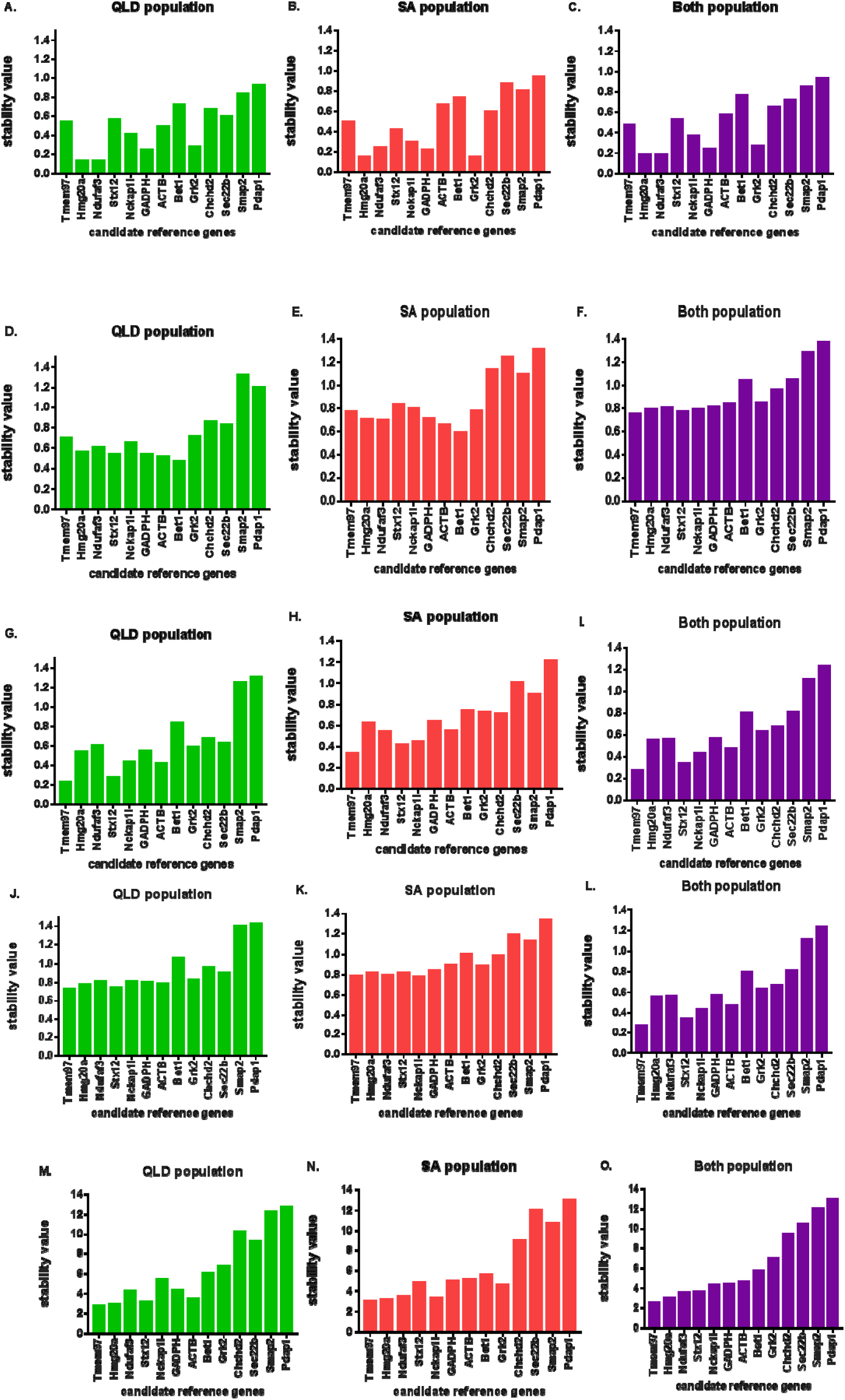
Evaluation of the stability of expression for 13 candidate reference genes in koalas using GeNorm (A-C), BestKeeper (D-F), NormFinder (G-l), the comparative ΔCt method (J-L) and the RefFinder tool (M-O). QLD represents Queensland population evaluation result; SA represents South Australian population; Both represents combination of the QLD and SA populations. The most stable reference genes have the lowest expression stability values.

To determine the optimal number of genes needed for RT-qPCR normalization, the average pairwise variation (V) was calculated between two consecutive normalization factors NF_n_ and NF_n+1_ across individual QLD and SA koala groups and in both populations. Generally, an additional reference gene is included at each step until V drops below 0.15. Below this point, additional reference genes are unlikely to have a beneficial impact for further improvement of data normalization^11,32^. In our study, the mean pairwise variation for the expression of genes ranked 2 and 3 was < 0.15 and remained <0.15 for the addition of all other reference genes across both QLD and SA subgroups and the combined group, as shown in Fig. 3.

**Figure 3:**
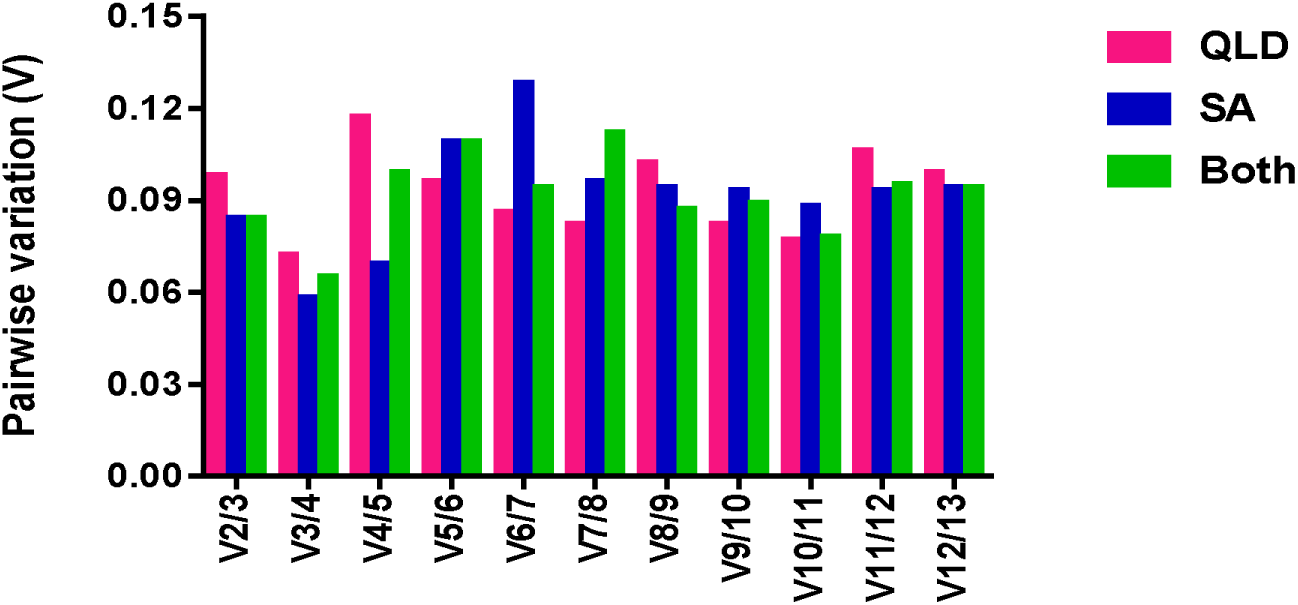
GeNorm calculation of pairwise variation analysis for choosing the optimal number of reference genes required for normalization, with a sequentially increasing number of genes. QLD: only QLD population evaluation result; SA: only SA population; Both: combined QLD and SA population. A value <0.15 indicates that the additional reference gene would not dramatically affect the normalization.

#### BestKeeper analysis

In the BestKeeper evaluation, *Bet1* and *ACTB* demonstrated the highest stability in QLD, SA and combined evaluation, a finding that was completely discordant with results from the other four algorithms. *Smap* and *Pdap1* exceeded the standard deviation (SD) threshold value of 1 across all geographical groups of this study and thus they were regarded as not suitable for subsequent gene expression normalization (Fig. 2 D-F). *Sec22b* and *Pdap1* also crossed the SD threshold value in the SA population and in the combined group. Overall, using this analytical approach, *Bet1, ACTB, Stx12, GAPDH, Hmg20a, Ndufaf3, Tmem97, Grk2* and *Nckap1l* remained stable in all sample sets.

#### NormFinder analysis

Based on NormFinder, the top five ranking candidate reference genes for QLD koalas were *Tmem97 > Stx12 > ACTB > Nckap1l > Hmg20a*. For SA animals, they were *Tmem97* > *Stx12 > Nckap1l > Ndufaf3 > ACTB*. For a combination of both populations, they were *Tmem97 > Stx12 > Nckap1l > ACTB > Hmg20a*. As shown in Fig. 2 G-l, Norm Finder suggested *Tmem97* was the most stable and *Stx12*, the second most stable gene across all experimental groups, which was in contrast with the geNorm output. Conversely, in alignment with geNorm and BestKeeper results, Pdap1 was identified as the least stable gene in QLD, SA and in the combined groups.

#### The comparative ΔCt analysis

According to the comparative ΔCt method, *Tmem97* was selected as most stable and *Stx12* was the second most stable gene in the QLD group and in combined group analysis. These results were very similar to NormFinder, but contrasted with results generated by the geNorm and BestKeeper algorithms (Fig.2 J-L). In contrast, *Nckap1l* was the most stable gene for the SA population and *Tmem97* was the second most stable. *Pdap1* was the most variable gene across all groups, this finding was consistent across all tested algorithms.

#### RefFinder evaluation

The four different algorithms produced divergent ranking of the stable genes. This variation has been noted in previous studies^33^. Based on the geometric mean value of each gene, comprehensive stability was evaluated through the RefFinder tool and candidate reference genes were ranked from 1 (most stable) to 13 (least stable) (Fig. 2 M-O). For QLD animals, the comprehensive ranking of candidate reference genes from 1 to 5 was *Tmem97* > *Hmg20a > Stx12 > ACTB > Ndufaf3*. For SA animals, it was *Tmem97* > *Hmg20a* > *Stx12 > Nckap1l > Ndufaf3*. For the combination of both populations, it was *Tmem97* > *Hmg20a > Ndufaf3> Stx12 > Nckap1l*. Overall, RefFinder chose *Tmem97* and *Hmg20a*, as the most stable genes in all of the samples as these expressed the lowest Geomean of the ranking values.

## DISCUSSION

Quantitative real time PCR has been used widely in molecular biology research to evaluate gene expression under certain conditions. To deliver biologically relevant data, normalization of gene expression with a reference gene is crucial. However, the selection of endogenous control genes is critical as gene expression should be stable in any experimental conditions or any type of tissues. In reality, there is no single universal stable gene available which is consistent in all experimental conditions. Consequently, reference gene stability needs to be verified prior to RT-qPCR studies for each tissue type. This study represents the first analysis of this type of evaluation on koala lymph node tissue. The stability of 13 candidate reference genes was assessed in koala submandibular lymph nodes across northern and southern koala populations. As shown in Figure 1, not all of the 13 analysed candidate genes had similar patterns of expression, suggesting variation in transcript abundance.

In this study, four different statistical algorithms were used to calculate the expression stability of 13 candidate reference genes for QLD and SA koala lymph node tissues with the QLD and SA populations analysed separately and also in combination. The comparative ΔCt method^8,34^ and geNorm^8^ algorithm evaluated the stability through intragroup differences and mean pairwise variation. The BestKeeper calculates the standard deviation of Ct values to determine stable genes and also used intragroup variation^35^ whereas NormFinder used both intra- and inter group variation. The RefFinder algorithm is used widely to calculate the geometric mean of the four algorithms results^36^. The outcome from the various algorithms is often dissimilar due to different approaches used by algorithms to evaluate the stability.

For the QLD population, *Tmem97* and *Stx12* were selected as the most stable genes through the ΔCt method and NormFinder. *Bet1* was most highly ranked by BestKeeper. *Hmg20a* and *Ndufaf3* were selected as the best pair of genes through geNorm pairwise variation evaluation. Regarding the SA population evaluation, ranking positions were dissimilar to the QLD population. In the SA population *Nckap1l* and *Tmem97* were selected as the most stable genes by comparative ΔCt method, while *Grk2* and *Hmg20a* were chosen as the best pair of stable genes through geNorm. The BestKeeper and NormFinder results for the two most stable genes in SA were similar to the QLD population ranking. In the combined population analysis, the ranking position of the two most stable genes was identical with the QLD population studies across different algorithms. Overall, the multiple algorithms produced dissimilar results, and RefFinder was used to solve this issue of discordance. In this comprehensive ranking, *Tmem97* and *Hmg20a* were selected as the two most stable genes for the subgroup and overall combination analysis of lymph node tissues.

*ACTB* and *GAPDH*, the most commonly used reference genes for normalization of quantitative expression studies in koalas^29,37,38^ were not ranked as the most stable genes with any of the algorithms or in any experimental group. The BestKeeper algorithm chose *ACTB* as the second most stable gene in all experimental conditions but in other algorithms and the final comprehensive analysis, its ranking position indicated lower stability. GAPDH was chosen as the third ranked reference gene based on geNorm algorithms, but its stability was lower with the other statistical algorithms and RefFinder.

In the geNorm algorithm evaluation, all reference genes expressed mean pairwise variation value of <0.15, indicating that there would be no improvement in the accuracy of data normalisation for the use of more than the top two ranked genes. However, in general three reference genes are recommended for accurate normalization in gene expression studies. Based on this study the top ranked genes overall in the RefFinder analysis in all conditions (*Tmem 97* and *Hmg20a*) would be good choices, potentially in combination with *Stx12* or *Ndufaf3* (in the top 5 ranked candidates for all groups analysed). *Tmem97* and *Hmg20a* are rarely used as endogenous controls to normalize expression levels in any species and there is no literature on their use in koala studies. *Tmem97*, a conserved integral membrane protein coding gene, is associated with cholesterol level maintenance^39^ and Hmg20a is involved with neuronal differentiation^40^ in mice. Transcripts of both were abundant in this study (Figure 1). Consequently, *Tmem97* and *Hmg20a* should be used in future normalisation studies on lymph node tissue in koalas. In addition, there may be further suitable reference genes that have not yet been explored. The use of the NGS generated transcriptomics data available for lymph node tissue to pre-select candidate genes with little variation in RNA expression for RT-qPCR normalisation appears to have been highly successful in that all the genes selected this way would be suitable for use in future studies. With essential tools now in place for marsupial gene normalisation (at least in koala lymph node tissue) from this study, future studies on relative gene expression and pathogen abundance now have a more robust method available for comparative gene expression RT-qPCR studies. Given that lymphocytes are the likely target cell for KoRV, a prevalent pathogen of koalas, lymph node expression studies are vital to understand whether KoRV can induce neoplasia or immunosuppression in koalas.

## METHODS

### Sample collection

Submandibular lymph nodes were collected from 19 wild adult koalas from northern (n= 11, South-East Queensland) and southern (n = 8, Lofty Ranges, South Australia) populations. The animals were euthanized due to sickness or injury of various causes (full details are in supplementary information Table SI). Prior to euthanasia, koalas were anaesthetised with 0.25 ml Zoletil (tiletamine/zolazepam) (Virbac) intramuscularly. Euthanasia was performed with an intravenous injection of lethabarb (pentobarbital) (Virbac). Tissues were collected within 3 hours of euthanasia, placed in RNA later (Qiagen) and stored at −80 °C until RNA extraction. All procedures were approved by the University of Queensland Animal Ethics Committee (Animal ethics number ANFRA/SVS/461/12 and ANRFA/SVS/445/15), Queensland Government Department of Environment and Heritage Protection (Scientific Purpose Permit WISP11989112), University of Adelaide Animal Ethics Committee (Animal ethics number S-2013-198) and SA Government Department of Environment, Water and Natural Resources (Scientific Purpose Permit Y26054).

### RNA extraction and cDNA synthesis

Total RNA was extracted using Qiagen RNeasy Plus Universal mini kit (Qiagen, Hilden, Germany) following the manufacturers instructions. RNA was further purified using an RNeasy mini kit (Qiagen, Hilden, Germany) with on-column DNase digestion (Qiagen, Hilden, Germany) as per the manufacturers guidelines. The quantity and quality of extracted RNA was assessed using a Nanodrop 1000 spectrophotometer (Thermo Scientific, Australia) and Agilent 2100 Bioanalyzer. cDNA was synthesized from 200 ng RNA using the QuantiTect Reverse Transcription kit (Qiagen, Germany) with a manufacturer provided oligo (dT) primer in a final volume 20 μl. The cDNA was diluted five-fold with RNAase free water and stored at −20 °C until required.

### Selection of candidate reference genes

These animals are a subset of those included in a larger transcriptomics study described elsewhere (Tarlinton et al., in preparation). Data from that study (available at the European Nucleotide Archive with the accession number PRJEB21505 assembled transcriptome at https://doi.org/10.6084/m9.figshare.5492512) were accessed for this study. Normalised abundances (TPM) for annotated genes were determined using Stringtie^41^ following mapping of reads using Hisat2^42^. Mean expression and variation across all replicates was determined and ranked for a) higher level of expression and b) low variation across all samples. The list of candidate stable genes is presented in Supplementary dataset 1 with their identifiers in the above data set, the sequences of the RNA transcripts of the selected genes are presented in Supplementary data set 2. A selection of the top 25 of these genes was then chosen for analysis of suitability as RT-qPCR normalisation genes via the MIQE guidelines. In addition to the stable genes identified from the transcriptome data, *ACTB* and *GAPDH* were also studied as these are the most commonly used reference genes in multiple koala studies.

### Primer design and amplification efficiency analysis for RT-qPCR

Gene specific primers for RT-qPCR were designed using the NCBI Primer-BLAST tool (https://www.ncbi.nlm.nih.gov/tools/primer-blast/). The assay was performed using Rotor Gene Corbett 6000 quantitative real-time PCR system (Qiagen). The size of the PCR products was between 95-145 bp and annealing temperature was optimized to 60 °C. cDNA was amplified using the PowerUp SYBR Green Master mix (ThermoFisher Scientific) following manufacturer instructions with a final reaction volume of 20 μl in each well containing 4 μl cDNA, 10 μl SYBR Green Master mix, 1 μl (10 μM) of each sense and anti-sense primer and 4 μl PCR-grade water. The PCR reaction was carried out with a hold temperature of 50 °C for 2 mins and 95 °C for 2 mins followed by 40 cycles of 95 °C for 15 s, 60 °C for 15 s and 72 °C for 1 min. To increase the accuracy and produce reliable results, all qPCR analyses were conducted with three replicates. For each primer pair, amplifications included a no-template control to ensure the absence of other contamination or primer dimer. Melting curve analysis was performed at the end of each PCR to verify primer specificity. Amplicons from each primer pair were tested by 2% agarose gel electrophoresis to verify the products’ size and absence of non-specific bands. The qPCR efficiency of each gene was determined by using slope analysis with a linear regression model. Undiluted cDNA samples were used to calculate the PCR efficiency and correlation coefficient (R^2^) for each primer pair based on the standard curve method. The standard curve was generated with five-fold serial dilutions of cDNA. The corresponding qPCR efficiencies (E) were calculated according to the equation E (%) = (10^(−1/slope)^ − 1) × 100^43^. Amplicons from the newly designed primers were Sanger sequenced and aligned with transcriptome data to confirm identity of the product.

### Determination and validation of expression stability of reference genes

Ct values for all samples were exported into an excel spreadsheet using Rotor-Gene Q 2.3.1.49 software. The average Ct values of three replicates were used for further analysis. The reference genes expression stability was analysed with commonly used statistical applets geNorm^8^, NormFinder^44^, BestKeeper^35^ and the comparative?ΔCt method^34^.

The GeNorm algorithm evaluates the expression stability value (M) of each housekeeping gene based on mean pairwise variation between selected candidate reference genes. The average ratio of two best stable genes expression levels should remain constant across all samples. The most stable gene shows the lowest M value and ultimately reference genes are ranked through stepwise elimination of genes with the highest M value. This statistical algorithm also evaluates the pairwise variation (V_n_/V_n+1_) to identify the optimal number of genes required for accurate normalization of real time PCR data^8^.

NormFinder determines expression stability through assessment of intra- and intergroup variation employing an ANOVA based approach. Genes with lower stability values exhibit higher expression stability in this algorithm^44^.

BestKeeper is used as an index to rank the stable reference genes through Pearson’s correlation coefficient as well as standard deviation (SD) and percentage covariance (CV) calculation of average Ct values. Analysed genes with SD values below 1 are considered as unstable and conversely in Pearson coefficients of correlation (R) analysis, the most stable genes exhibit values closest to 1^35^.

The comparative ΔCt method compares the relative expression value of possible reference genes in pairs within each studied sample and ranks based on reproducibility of gene expression variation among experimental samples^34^.

Each algorithm might rank the candidate reference genes differently. To solve this issue, the final reference gene ranking was conducted using RefFinder (http://150.216.56.64/referencegene.php), a web based comprehensive evaluation tool (Cotton EST database, RefFinder software) that assesses all statistical algorithms and ranks the stable genes based on the geometric mean values. Analyses were done in three separate groupings, (a) Queensland koalas (QLD, northern population) (b) South Australian koalas (SA, southern population) and (c) combination of both groups.

## ACKNOWLEDGMENT

This project and scholarship for NS were funded by the Queensland Department of the Environment and Heritage Koala Research Grant Programme 2012. NS was also supported by a Keith Mackie Lucas travel scholarship from the University of Queensland. Koalas for post mortem were accessed through the Moggill Wildlife hospital (QLD Department of the Environment and Heritage Protection) and Fauna Rescue of South Australia and we are extremely grateful for the staff and volunteers that work with these organisations for their work in koala rescue.

## AUTHOR CONTRIBUTIONS

N.S. performed RNA extraction, primer designing, RT-qPCR, laboratory experiments, data analysis and drafted manuscript. R.D.E and J. Moreton performed bioinformatics analysis and edited the manuscript. J.F. and N. Speight helped in sample collection and reviewing manuscript. F.H, D.T. and L.W. reviewed the manuscript. J. Meers, J.M.S., G.S. and H.O. helped in laboratory experiment set up, data interpretation and manuscript preparation. R.E.T initiates the study design, coordination of work and edited the manuscript. All authors read and approved the final manuscript.

